# SEMITONES: Single-cEll Marker IdentificaTiON by Enrichment Scoring

**DOI:** 10.1101/2020.11.17.386664

**Authors:** Anna Hendrika Cornelia Vlot, Setareh Maghsudi, Uwe Ohler

## Abstract

Identification of markers is an essential step in single-cell analytic. Current marker identification strategies typically rely on cluster assignments of cells. Cluster assignment, in particular of development data, is non-trivial, potentially arbitrary and commonly relies on prior knowledge. Yet, cluster uncertainty is not commonly taken into account. In response, we present SEMITONES, a principled method for cluster-free marker identification. We showcase its application on healthy haematopoiesis data as 1) a robust alternative to highly variable gene selection, 2) for marker gene and regulatory region identification, and 3) for the construction of co-enrichment networks that reveal regulators of cell identity.

## Background

Since the inception of single-cell RNA sequencing (scRNA-seq) in 2009, singlecell methods have become commonplace. scRNA-seq provides a snapshot of the gene expression state of single cells and is a valuable resource to address questions on cell identity and cell lineage relationships. In recent years, single-cell assays for transposase-accessible chromatin using sequencing methods (scATAC-seq) have also become available. scATAC-seq provides a snapshot of the chromatin accessibility profile of single cells and can be used to identify putative cell-type-specific *cis*-regulatory regions.

The appearance of these novel, sparse data types sparked the development of specialized single-cell analysis methods that cover the entire single-cell data analysis workflow. In both scRNA-seq and scATAC-seq pipelines, feature identification is an essential step which is commonly performed twice. First for feature selection to reduce the number of genes or accessible regions in the data, and later to identify markers of cell identity [1]. Feature selection for dimensionality reduction is most commonly performed by the identification of a certain number of the most variable or most common features. The number of features depends on the task complexity and influences clustering accuracy [1, 2]. If too many features are chosen, spurious clusters of cells with no specific identity may occur. Contrarily, if too few genes are selected, clusters of cells from distinct biological origins may cluster together. This is especially problematic since the ground truth of the cell types present in an experiment is commonly not available. Additionally, these effects are propagated into the downstream analyses including marker identification, where commonly performed differential expression methods rely on the premise that the cell identities are known without consideration for annotation uncertainty [3]. These observations illustrate the interdependence between clustering accuracy and feature identification.

The aforementioned difficulties are aggravated when one considers developmental data, like whole-organism development [4] or haematopoiesis [5] data. In these datasets, cells are found along the full developmental axis, from omni- and pluripotent stem cells to fully differentiated cells. Thus, clustering the cells into distinct cell types becomes less meaningful. Pseudotime analysis is common for data of this nature. In these analyses, marker feature identification is commonly performed by differential testing between branches, without considering the uncertainty in branching point determination. Thus, reservations considering annotation accuracy persist.

Finally, genes and *cis*-regulatory elements act in interaction with one another. It is the combination of expressed genes and/or open chromatin regions which determine the transcriptomic or *cis*-regulatory state of an individual cell. Thus, the identification of distinct regulatory (gene expression) networks is expected to provide a clearer picture of cell identity than individual markers.

To address the aforementioned challenges, we have developed SEMITONES (Single-cEll Marker IdentificaTiON by Enrichment Scoring). SEMITONES is a method for the identification of informative features and/or feature sets in scRNA-seq and scATAC-seq data independent of data clustering. We illustrate the practical use of SEMITONES by application to published healthy haematopoiesis scRNA-seq and scATAC-seq data [5]. We show its application to feature selection for dimensionality reduction, marker gene and cis-regulatory element identification, and signature gene set identification. In short, we present a flexible method for the identification of signatures of cell identity in single-cell omics data.

## Results

SEMITONES identifies informative features in single-cell omics data. We consider a feature as informative if it is only present or absent in similar cells since both the presence and absence of a given feature are informative for cell identity. The standard SEMITONES workflow consists of three steps. First, it selects a set of diverse reference cells from the entire population of cells to serve as a representation of the cell states present in the sample (Figure 1a). Next, it calculates the enrichment score of each feature for the reference cell neighbourhood using a linear regression framework (Figure 1b). Lastly, to decide whether a future is informative or not, it performs statistical testing against a permutation null distribution (Figure 1c). Besides single features, this procedure can be followed for sets of features. The feature set enrichment scores can then be used to construct co-enrichment graphs where vertices represent features and the edges between them are weighted by enrichment scores (Figure 1d).

**Figure 1:**
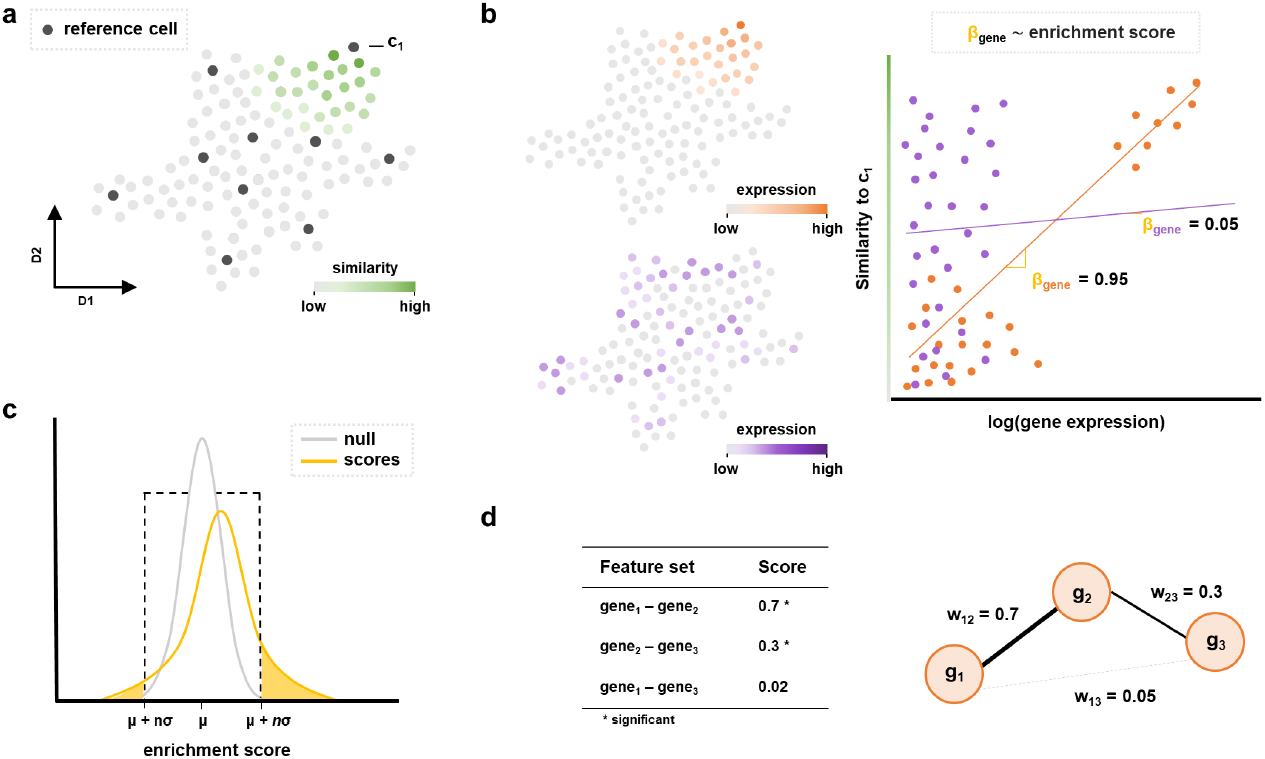
SEMITONES workflow. a) a two-dimensional embedding of all cells where dark grey dots are the selected reference cells. In the green gradient, we show the similarity to reference cell c_1_. b) Based on the assumption that informative genes are only expressed in the reference cell neighbourhood, we identify informative (orange) and uninformative (purple) genes in the reference cell (c_1_) neighbourhood. The value of β is (proportional to) the enrichment score, so informative genes get high scores and vice versa. c) Scores in the shaded orange area are declared significant because they are more than n standard deviations away from the mean of the null-distribution. This null-distribution is the distribution of enrichment scores for the permuted feature vector of all features in the data. d) Given enrichment scores for sets of genes, we construct co-enrichment graphs where vertices are genes and the edges are weighted by the enrichment score of the gene set consisting of the genes connected by this edge.

We evaluate the application of SEMITONES on published scRNA-seq and scATAC-seq data of healthy haematopoiesis [5]. The primary objective of SEMI-TONES, the clustering-free identification of markers of cell identity by enrichment scoring, is explored for both scRNA-seq and scATAC-seq. Additionally, we explore the selection of significantly enriched genes for feature selection as an alternative to the selection of highly variable genes. Lastly, we show how SEMITONES can be used to construct co-enrichment networks which reveal regulators of cell identity.

### SEMITONES identifies marker genes

SEMITONES identifies known marker genes without preceding clustering (Figure 2). To illustrate, the top 3 most highly enriched genes include the erythrocyte markers *AHSP, HBB*, and *CA1* [6], the plasma cell markers *TNFRSF17* [7] and *GPRC5D* [8], and the eosinophil/basophil/mast cell markers *HDC* and *CLC* [9] (Figure 2a, see Supplementary Table 1, Additional File 2). This confirms that SEMITONES identifies markers of specialized cell types. In addition, SEMITONES identifies markers for stem- and progenitor cells, like the haematopoietic stem cell (HSC) markers *AVP* [10] and *CRHBP* [11], the HSC and multipotent progenitor (MPP) marker *SPINK2* [12], and the transcription factor *GATA2* associated with erythroid-megakaryocyte lineage commitment [6] (Figure 1b, see Supplementary Table 1, Additional File 2). SEMITONES can also identify markers for specialized subpopulations of highly similar cells, including the CD4^+^ T helper 17 (T_h_17) cell marker *TNFRSF4* [9], the CD8^+^ mucosal associated invariant T (MAIT) cell marker *SLC4A10* [13], and the transitional B cell specific *DTX1* (Figure 2b, see Supplementary Table 1, Additional File 2). These results illustrate that SEMITONES identifies markers of cell identity-specific marker genes for fully differentiated, progenitor, and rare cell populations.

**Figure 2:**
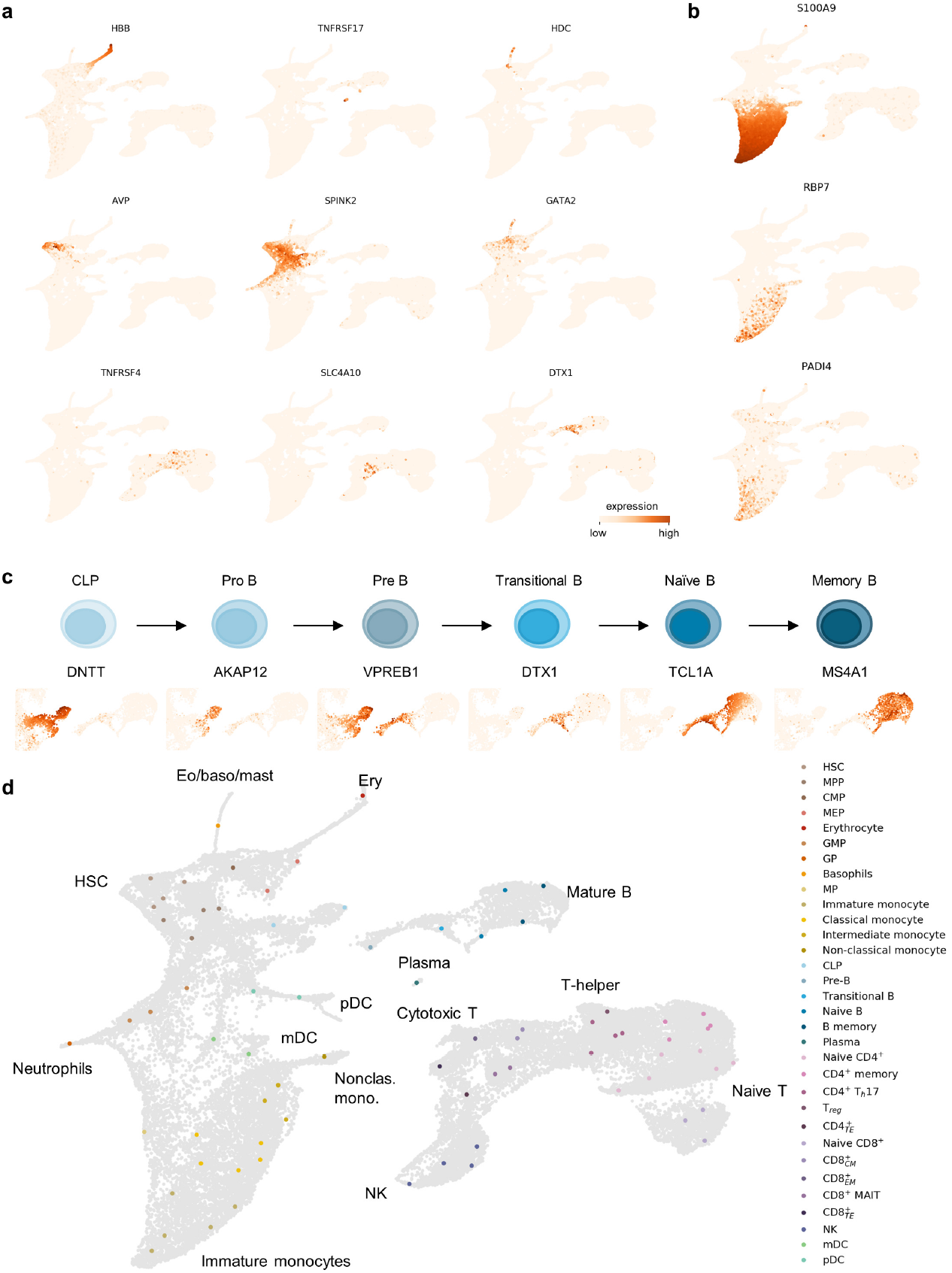
Application of SEMITONES for marker gene identification in scRNA-seq data. a) Highly specific markers of well-characterized cell types (top row: erythrocytes, plasma cells, eosinophil/basophil/mast cell lineage), progenitor cells (middle row: haematopoietic stem cells, haematopoietic stem- and progenitor cells, myeloid progenitors), and specific subpopulations (bottom row: T_reg_, CD8^+^ MAIT, and transitional B cells). b) The expression profile of the known immature monocyte marker *S100A9* and the newly proposed immature classical monocyte markers *RBP7* and *PAD41*. c) The expression of markers along the B cell developmental trajectory. d) Reference cell annotations based on the marker genes identified by SEMITONES.

SEMITONES is also suited to retrieve markers of specialized subpopulations of highly similar cells, such as specific markers for different monocytic cell populations. To illustrate, SEMITONES identified relative enrichment markers that distinguish immature classical monocytes, classical monocytes, and intermediate monocytes (see supplementary Figure 1, Additional File 1, and Supplementary Table 1, Additional File 2). Here, immature classical monocytes were identified by top 4 enrichment of *S100A8, S100A9*, and *S100A12* and relatively lower enrichment of the classical monocyte markers *CD14* and *VCAN*. The *S100A9* and *S100A8* genes have previously been described to be highly expressed in the early stages of monocytic differentiation [14, 15, 16]. Additionally, these *S100* genes are also markers for human monocytic myeloid-derived suppressor cells (MDSCs) that develop from immature myeloid cells in disease states like chronic inflammation [17], further corroborating this annotation. Using SEMITONES we identify *PLBD1*, *RBP7*, and *PADI4* as highly enriched in immature classical monocytes (Figure 1c, see Supplementary Table 1, Additional File 2). These three genes are not within the top 10 most highly enriched genes for other monocytic subpopulations, and the co-expression of *RBP7* and *PADI4* appears to be specific to immature classical monocytes (Figure 2b). Similarly, we identify reference cells with high enrichment for *LGALS2* in absence of top 10 enrichment of the classical monocyte marker *VCAN* [9]. In line with observations of higher relative expression of *LGALS2* in intermediate monocytes compared to non-classical monocytes [18], we suggest that this identifies a population of intermediate monocytes.

Similarly, SEMITONES identifies markers of developmental stages along the haematopoiesis axis, as we illustrate using the example of B-cell maturation (Figure 2c). Here, as the top-scoring gene in one reference cell, we identify the *DNTT* gene which codes for the recombination substrate terminal deoxynucleotidyl transferase that is involved in immunoglobulin (Ig) and T-cell receptor (TCR) recombination [19]. This gene is expressed in the lymphoid-primed progenitor (LMPP) stage and upregulated in the common lymphoid progenitor (CLP) stage [20]. Therefore, we can identify this cell as a CLP. In another cell for which high *DNTT* enrichment is found, the top-scoring enrichment is found for *AKAP12* (see Supplementary Table 1, Additional File 2), which is expressed exclusively in pro-, pre-, and immature B lymphocytes [21]. Given the combined enrichment of *DNTT* and *AKAP12*, we identify this cell as a pro-B cell. Both these cells also show enrichment for the *VPREB1* gene, which encodes the ι polypeptide chain that is part of the pre-B cell receptor [22]. This gene is lowly expressed in CLPs and highly expressed in pro-B and pre-B cells [23], further confirming our annotations. Interestingly, the identification of a cell stage with a strong cell cycle signature which includes the *TOP2A, KIFC1*, and *NUSAP1* genes, alongside *VPREB1* as the 19^th^ most enriched gene, allows for the identification of large-pre B cells, a highly proliferative cell state in B cell development [24]. Furthermore, we find high *DTX1* and *BMP3* enrichment for a cell that can now be annotated as a transitional B lymphocyte, the next step in B lymphocyte development [25, 26]. Next, selective top 10 enrichment of *TCL1A*, which is not expressed in memory B cells [27], and *FCER2*, which is involved in B cell differentiation and regulates IgE production [22], indicates cells that are immature B lymphocytes [9]. Lastly, top enrichment for *MS4A1*, coding for the B-lymphocyte antigen CD20 which promotes calcium influx after activation by the BCR [28], and *FCER2* in the absence of *TCL1A* can be used to identify mature B cells [9, 27]. To conclude, SEMITONES identifies markers of many cell identities along the developmental axis without the need to enforce (arbitrary) cell identity boundaries.

After confirming that SEMITONES identifies markers of cell identity, we use the top 10 most highly enriched genes to annotate all reference cells (Figure 2d). To evaluate the cell-type retrieval of our data-driven selection approach, we compare those annotations to the published cluster annotations from [5]. This comparison reveals that one cell of every annotation is present in our set of 75 reference cells (see supplementary Figure 2, Additional File 1), i.e., our simple data-driven selection procedure manages to include all cell types of interest by selecting just 0.2% of the total number of cells as reference cells. Besides, we identify additional cell types based on SEMITONES reference cell selection and enrichment scores, including intermediate monocytes, and several B- and T-lymphocyte subsets. Further comparisons were made to a set of 75 manually selected reference cells (see supplementary Figure 3, Additional File 1), annotated based on SEMITONES enrichment scores. One of these manually selected reference cells was identified to be a plasmablast, a cell type that is not part of the cluster-based or algorithmically selected reference cell annotations. We note that the data-driven reference cell selection depends on the dissimilarity metric and the embedding over which the dissimilarity is determined (see supplementary Figure 4, Additional File 1). In general, given a descriptive similarity metric, the data-driven selection of reference cells will provide a sample of cells that is representative of the population.

The results described above relate to enrichment scores obtained using an RBF-kernel with γ = 8 × 10^−1^ to represent the pairwise cell similarities because this parameterization allows for the identification of selective cell identity markers. However, by decreasing the value of γ, one can also identify more globally enriched genes (see supplementary Figure 5, Additional File 1). Namely, γ is the inverse of the radius of influence, which is proportional to the size of the cell neighbourhood for which we want to retrieve marker genes. This illustrates how SEMITONES can flexibly infer highly specific or global cell identity markers without relying on hard cluster boundaries.

### SEMITONES identifies transcriptional regulators

To reveal co-enrichment relationships of genes in a given cell neighbourhood, we construct co-enrichment graphs using SEMITONES co-enrichment scores. Since ~ 143 × 10^6^ possible pairwise gene sets of 16,900 expressed genes exist, we compute pairwise enrichment scores for gene sets of significantly enriched genes in a subset of reference cells. This subset of reference cells contains one cell of each annotation, where we select the cell with the enrichment score for the primary annotation marker (see Supplementary Table 2, Additional File 3). Given this subset, we obtain 333974 possible pairwise sets of significantly enriched genes (n_σ_ = 25) per cell in the subset. Next, we perform enrichment scoring for all gene sets for each reference cell in the subset (see Methods). We then construct co-enrichment graphs containing all gene sets that are significantly, positively co-enriched (n_σ_ = 30) in some reference cell. To unveil the crucial connections in each co-enrichment graph, we evaluate the maximum spanning tree (MST) of each graph.

The co-enrichment graphs contain paths that link together genes that interact in specific stages of haematopoietic development. To illustrate, in the interaction co-enrichment graph of the transitional B cell neighbourhood we find that *IGLL5* is highly connected to its predicted interaction partner*CD79B* (see supplementary Figure 6, Additional File 1, [29]). These genes encode proteins (Igβ, and Igλ, respectively) that are involved in pre-B cell receptor signalling, which is an essential process in the development of immature B cells [30]. Another example concerns *CD3E, CD3D* and *CD8A*, for which interactions are found in the co-enrichment graph of naive CD8^+^ (Figure 3a), which are predicted in the STRING database [29] and have a mechanistic basis. Namely, the T-cell surface glycoprotein CD8 is thought to play a major role in the targeted delivery of the Lck protein to the CD3-complex, of which the CD3ε chain and CD3δ chain are part, during T cell activation [31]. These results illustrate that SEMITONES can identify biologically meaningful and cell identity specific co-expression graphs from scRNA-seq data.

**Figure 3:**
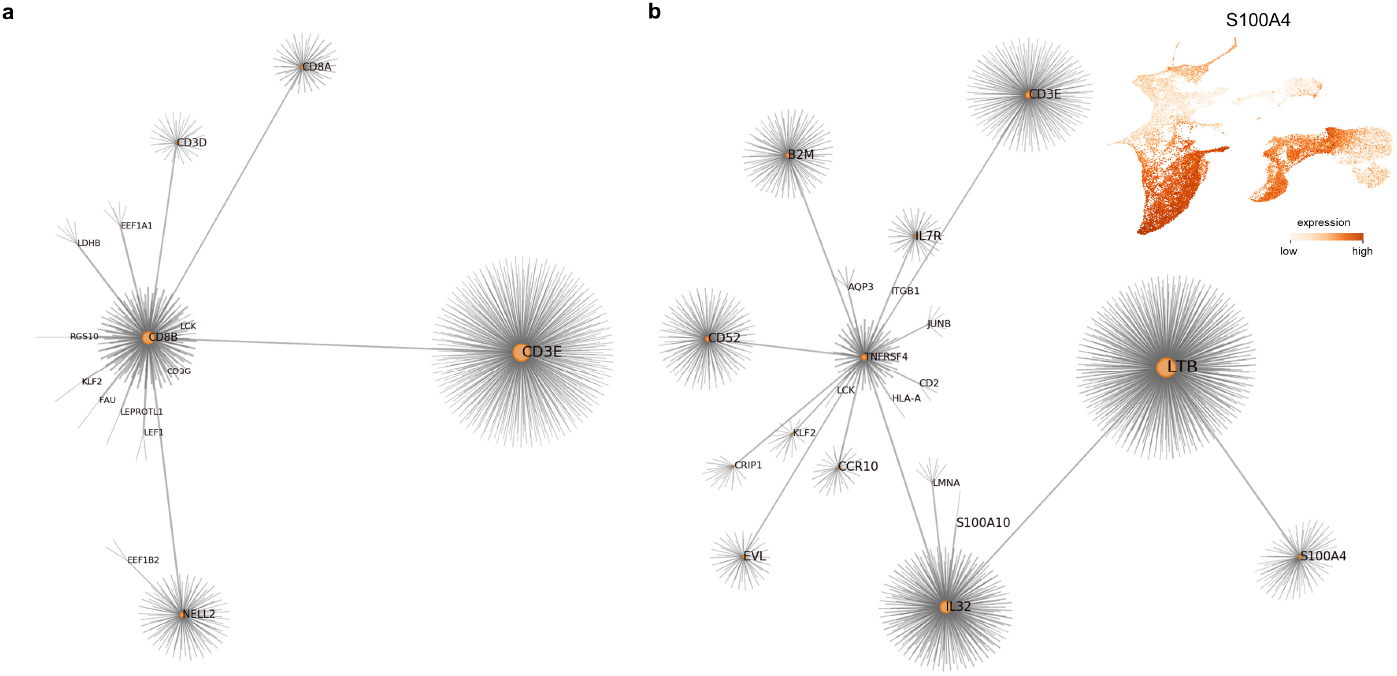
Gene co-enrichment graphs constructed from SEMITONES gene set enrichment scores. a) The maximum spanning tree (MST) of the interaction co-enrichment graph for naive CD8^+^ cells. b) The interaction co-enrichment graph of T_h_17 cells shows a central role for the *S100A4* gene which is not selectively expressed in regulatory T-cells. In both graphs, the vertex size is proportional to their weighted degree.

Inspection of the interaction co-enrichment graphs for the monocytic cell populations reveals subtle differences in otherwise highly similar graphs. For example, a central role for *S100* genes are found for all populations. Notably, the *S100A9* gene plays a more central role in the co-enrichment graph of the classical monocytes than that of the immature classical monocytes (see supplementary Figure 7, Additional File 1), whilst the *S100A9* gene was found to be more highly enriched in immature classical monocytes. Differences between monocytic cell populations are also found regarding which other genes a highly connected gene connects to. To illustrate, *NEAT1* is a highly central node in classical and intermediate monocytes but connects to different genes in each subset (see supplementary Figure 7, Additional File 1). In classical monocytes, *NEAT1* is highly connected to *S100A9* and *S100A8*, but in intermediate monocytes it is highly connected to *LGALS2, HLA-DRB1*, and *HLA-DRA*. These interactions are plausible based on higher expression of *S100A8* and *S100A9* in classical monocytes and *HLA-DRA* in intermediate monocytes compared to other monocyte populations [32]. Descriptions of gene interactions in monocytic subpopulations are only sparingly available in the literature, and we did not find any mechanistic evidence for the interactions described above. In general, however, these results confirm that SEMITONES uncovers subtle differences in the co-enrichment networks of highly similar cell populations. Based on the evidence described for transitional B lymphocytes and naive CD8^+^ cells, we suggest that these represent putative mechanistic distinctions.

To identify central genes in the co-enrichment networks, we calculate the current flow betweenness centrality of the vertices in the MST (see Supplementary Table 3, Additional File 4). Inspection of the top 10 most central genes reveals known regulators of cell identities. For example, the known regulators of erythropoiesis *GFI1B* and *HES6* are in the most central vertices of all erythrocyte co-enrichment graphs [33, 34]. Interestingly, these genes are ranked only 31st and 46th most enriched in the erythrocyte neighbourhood. Similarly, *S100A4* is a highly central node in all T_h_17 co-enrichment graphs (Figure 3b, see Supplementary Table 3, Additional File 4), but is only the 52nd most enriched gene in the T_h_17 neighbourhood. This rank is intuitively coherent with the global expression profile of this gene, with high expression observed in all monocyte, monocytic dendritic cells (mDC), natural killer (NK) cell, and mature T lymphocyte populations (Figure 3b). *S100A4* is suggested as a regulator of T_h_17 differentiation, albeit in Rheumatoid Arthritis mouse models [35]. Another *S100* gene, *S100A11*, was found as a highly central node in minimum expression-based co-enrichment graphs of T_h_17 and T_reg_ neighbourhoods (see Supplementary Table 3, Additional File 4), whilst ranking as 546th and 790th most enriched, respectively. This gene has been implicated as a regulator of T_reg_ development [36]. Based on these results, SEMITONES co-enrichment analysis enables the identification of putative regulators of cell-type specialization, also in cases where these regulators are not restrictively expressed in a specific cell neighbourhood.

Lastly, we qualitatively assess the robustness of biologically meaningful co-enrichment identification when using different approaches for gene set expression vectors. We find that some, but not all, connections are shared between all graphs for a given cell neighbourhood. For example, *AHSP* and *GFI1B* are found to be highly connected in all erythrocyte co-enrichment graphs (see supplementary Figure 8, Additional File 1abcd), whilst the connections of *AHSP, HBB*, and *HBB* are only found for interaction, maximum value-, and median value-based co-enrichment graphs (see supplementary Figure 8, Additional File 1abc). Similarly, the interactions between *ASHP, KLF1, HBA1* and *HBA2* are only found in the minimum-value based graphs (see supplementary Figure 8, Additional File 1d). All these connections can be traced back to the biology of erythrocyte development and function. Namely, both *AHSP* and *GFI1B* are essential for erythrocyte development and function [33, 37], AHSP is a haemoglobin stabilizing protein and a chaperone of *HBA1* and *HBA2*, and KLF1 binds to the promoters of *HBA1* and *HBA2* [38]. Additionally, high confidence interactions from experiments and curated databases were found between *AHSP, HBA1, HBA2, HBB*, and *HBD* in STRING [29]. Overall, these results suggest that interaction, maximum value, and medium value-based coenrichment are more similar to each other than the minimum expression-based co-enrichment graphs. This is readily explained by the relative focus on more lowly expressed genes for the minimum value-based co-enrichment graphs, with its stronger emphasis on lowly expressed genes. Importantly, independent of the method of gene set expression vector construction, biological proof of the co-enrichment identified SEMITONES can be found in curated databases and the scientific literature [29].

### SEMITONES for feature selection

SEMITONES is also a highly effective approach for feature selection. Feature selection typically takes place at the beginning of the preprocessing, when little information is available to aid the selection of a suitable similarity metric. We therefore choose the standard cosine similarity to characterize the similarity between cells. Additionally, we compute the similarity over the top 50 principal components instead of an optimized multi-dimensional UMAP. On the same grounds, we use the manually curated reference for feature selection. These reference cells were annotated when assessing the cell type retrieval in the data-driven reference cell selection, and we will use the same annotations in this section. We use SEMITONES to select 4000, 2000, 1000 and 500 significantly enriched genes by adjusting the number of standard deviations away from the null-distribution mean at which we declare significance. This approach is compared to selecting the same numbers of highly variable genes (HVGs) with the “Seurat-flavoured” HVG selection from Scanpy [39]. For each set of selected features, we perform LSI and construct a 2D UMAP using 35 nearest neigh-bours with a minimum Euclidean distance of 0.3. The methods are evaluated based on the cell identity separation and marker gene localization in the UMAP space.

The separation of cell identities in the UMAP space is less affected by selecting a lower number of SEMITONES enriched genes than to selecting a lower number of HVGs (see supplementary Figure 9, Additional File 1). To illustrate, when selecting 4000 genes, the only difference is the reduced separation of erythrocytes and megakaryocyte-erythrocyte progenitor (MEP) cells when using HVGs. However, the discrepancies increase as we lower the number of genes we select. For example, when using the top 2000 HVGs, plasmablasts and plasma cells cluster together, and one CD8^+^CM is found within the naive CD8^+^ cell neighbourhood. There is no decreased separation when using 2000 most enriched genes, and even lowering the number of genes to the 1000 most enriched genes does not impact the separation of cell identity in the 2D UMAP space. In contrast, when using the 1000 most variable genes, one B cell is found in the NK-cell neighbourhood, and one NK cell is found in the CD4^+^ neighbourhood. Additionally, granulocytes, plasmacytoid dendritic cells (pDCs), and granulocyte and monocyte progenitors (GMP) are no longer as distinctly separated, and neither are the different monocyte subpopulations (Figure 4). Lastly, and perhaps most apparent, the erythrocyte and eosinophil/basophil/mast cell populations are not resolved. Strikingly, these erythrocyte, eosinophil/basophil/mast cell and the granulocyte progenitor (moving towards neutrophils) branches that contain only a few cells, are still resolved when using just the 500 most highly enriched genes, but we observe the first loss of resolution, with the naive CD8^+^ cells merging with the CD4^+^ and T_h_17 cell neighbourhoods. The naive CD8^+^ population remains well separated when using the top 500 HVGs but at the cost of a further decrease in the separation of additional T cell subsets and monocyte subpopulations. Overall, these results demonstrate that a smaller number of SEMITONES selected features explains the same amount of biological variation as a larger set of HVGs.

**Figure 4:**
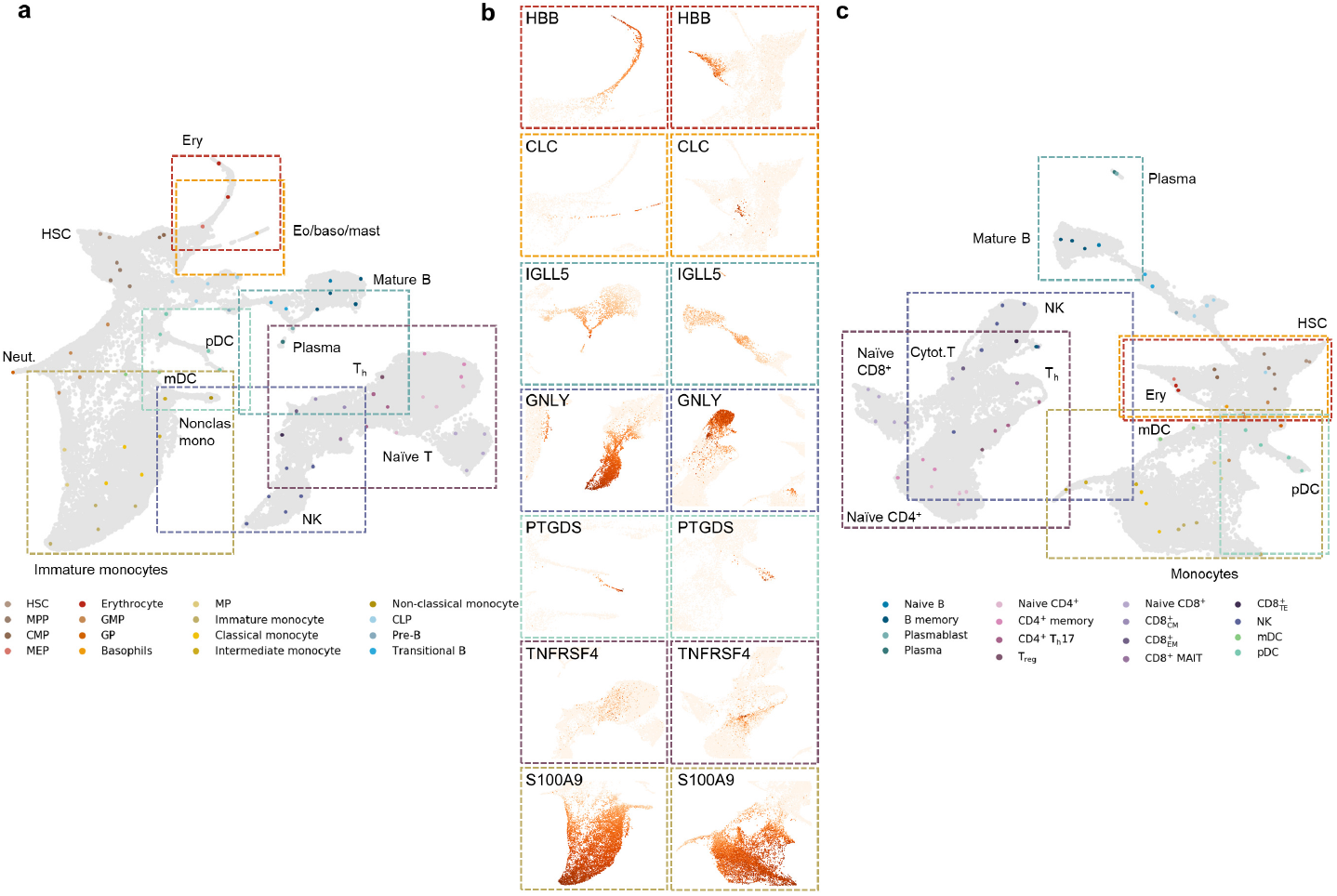
SEMITONES is a more sensitive alternative to highly variable gene selection. a) UMAP embedding of the scRNA-seq data based on the top 1000 most enriched genes. b) Marker gene expression visualized on the top enriched gene UMAP (left column, panel a) and the highly variable gene UMAP (right column, panel c). The border colours correspond to the annotation label colours of the label for which this gene is a marker. c) UMAP embedding of the scRNA-seq data based on the top 1000 most variable genes.

The decreased separation of cell identities when using HVGs compared to highly enriched genes can be linked to the absence of marker genes in the HVG sets: the erythrocyte markers *AHSP* and *HBB* are absent from the top 4000 HVGs onward, and the plasmablast marker *IGLL5* and the CD8^+^M and NK-cell expressed *CCL5* are absent from the top 2000 HVGs. Additionally, differences in the separation of cell identity can be linked to differences in marker gene localization in the UMAP space as illustrated in Figure 4 for 1000 enriched versus 1000 HVG genes. There are also genes that are more localized in UMAPs constructed using the top HVGs, e.g. the T_h_17 marker *TNFRSF4* when using 1000 (Figure 4) or 500 (see supplementary Figure 10, Additional File 1) genes. In general, however, gene expression appears more localized in UMAPs constructed using the top SEMITONES enriched genes. Overall, these results suggest that SEMITONES feature selection reduces the gene space to a small set of highly informative genes.

### SEMITONES identifies cell-specific cis-regulatory elements

Since SEMITONES is readily compatible with the scATAC-seq input matrices, we also explore its application for the identification of enriched ATAC peaks for 75 algorithmically selected reference cells. Visual inspection of top and bottom scoring regions reveals that SEMITONES accurately identifies peaks that are specifically present or absent in specific cell neighbourhoods (Figure 5a). Rarely, these significantly enriched peaks correspond to known *cis*-regulatory regions, like the PID1-DNER Intergenic CAGE-Defined Monocyte Enhancer (chromosome 2, 230147763-230148263bp, Figure 5a). Therefore, we use GREAT (v4.0.4) [40] to identify associated genes and enriched GO terms for the significantly enriched peaks (n_σ_ = 20). Based on this, we confirm that the peaks are in regions responsible for haematopoietic (e.g. HSC differentiation) and immune (e.g. leukocyte degranulation) processes. Many of these terms were cell typespecific and enabled us to directly annotate 74 out of 75 reference cells based on the GO terms and their associated genes (Figure 5b), without having to fall back on complementary data such as from scRNA-seq. Most of the annotations are concordant with the annotations in [5], which were obtained using Seurat’s canonical correlation analysis with scRNA-seq based annotations, with the exception of resolving all monocyte and T lymphocyte subpopulations. However, in turn, we reveal additional signatures, including a Notch signalling signature indicative of a pre-T lymphocyte fate in a CLP and a dendritic signature in a subset of reference cells. These observations may be related to the selection and representation of the reference cells and their neighbourhoods, and they support the notion that early *cis*-regulatory signatures reveal lineage commitments before they can be identified on the RNA level. Separate inference on the chromatin level is thus essential to gain novel insights from scATAC-seq data.

**Figure 5:**
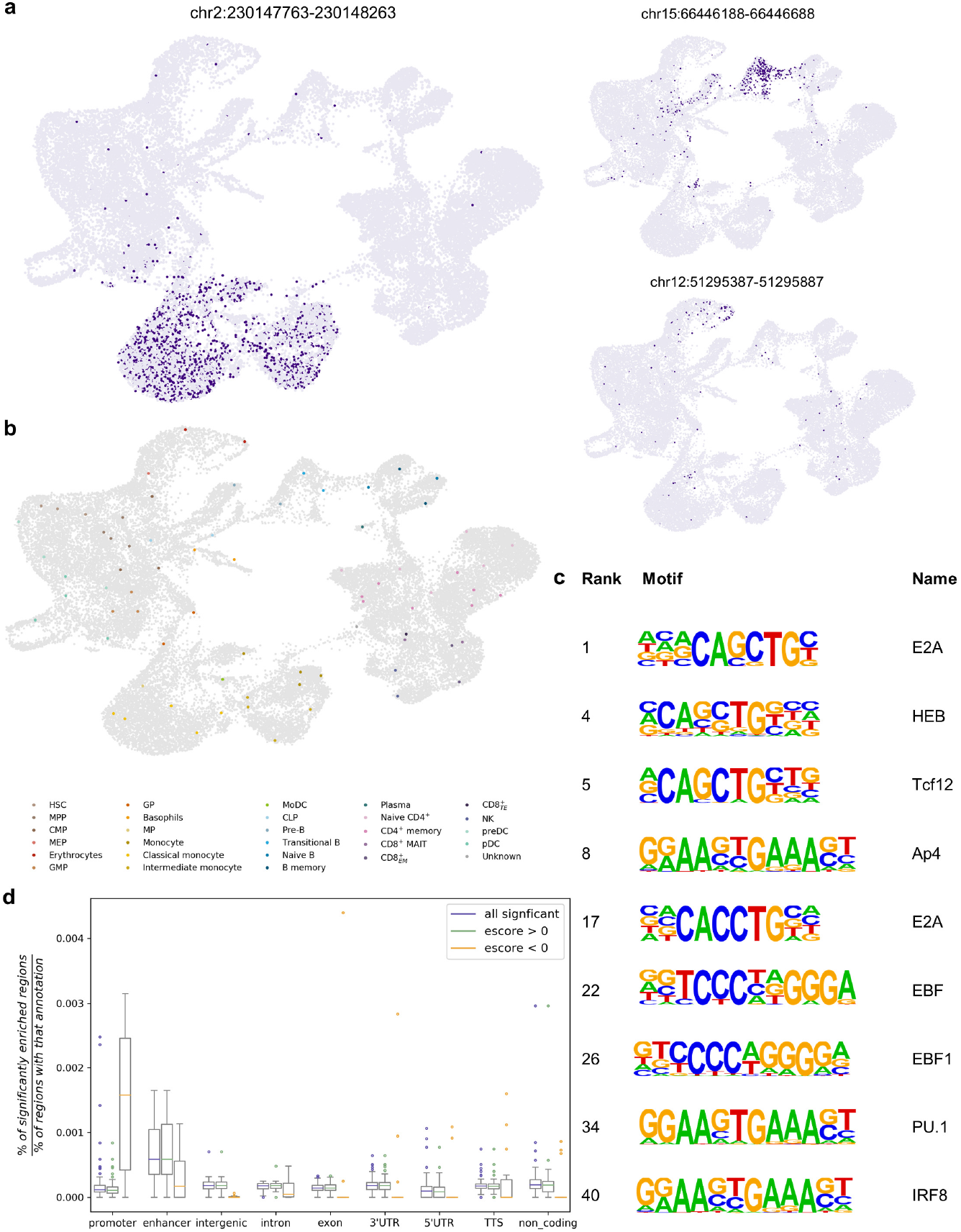
Application of SEMITONES for marker region identification in scATAC-seq data. a) The accessibility profile of several highly enriched regions visualized in a UMAP embedding. Region chr2:230147763-230148263 contains the PID1-DNER Intergenic CAGE-Defined Monocyte Enhancer. b) Reference cell annotations based on enriched GO-terms and associated genes, identified by HOMER, in significantly enriched marker regions identified by SEMITONES. c) Motifs that were found to be enriched in regions that SEMITONES identified as significantly enriched in transitional B cells. d) The normalized percentage of significantly enriched regions that have a certain annotation in HOMER or FANTOM5.

Significantly enriched peaks are enriched for the transcription factor binding motifs that one would expect to find in the annotated cell types. For example, HOX motifs are enriched in stem and progenitor cells, GATA motifs are enriched in the myeloid lineage, and PRDM1 and IRF4 motifs are enriched in the B cell lineages. We also identify cell type-specific motifs. For example, we find enrichment for motifs of the known regulators of B cell differentiation E2A, EBF, PAX5, PU.1 and IRF8 [41] in pre-B and transitional B cells (Figure 5c), and for GATA3, which is indispensable for T helper 2 (T_h_2) cell differentiation [42], in the CD4^+^ memory cell neighbourhood. These results further suggest that SEMITONES identifies distinct and import features (i.e. open chromatin regions) of cell identity.

Finally, we evaluate whether certain *cis*-regulatory elements are overrepresented in the significantly enriched peaks. Selectively inaccessible regions are more often promoter regions than any other *cis*-regulatory regions (Figure 5d). In the same vein, peaks with a positive enrichment score are, on average, most likely to fall in enhancer regions. Both these trends fit prior analyses that showed that in general, promoters per default are open across conditions, while many distal regulatory regions are specifically opened [43]. Lastly, we identify a relative overrepresentation of enhancer regions in monocytes and T lymphocytes (see supplementary Figure 11, Additional File 1), although this might be related to a relative overrepresentation of certain cell types in the FANTOM5 data. Overall, these results illustrate how the identification of cell type-specific open chromatin regions is invaluable to the elucidation of the role of *cis*-regulatory element accessibility in the acquisition and maintenance of cell identity.

### Scalability of SEMITONES

SEMITONES calculates enrichment scores of 30,000 features for a single reference cells in just a few minutes when the number of non-zero values is representative of scRNA-seq (~ 10% non-zero values) or scATAC-seq (~ 2% non-zero values, Figure 6bc). When applying SEMITONES to large and sparse data with a density representative of scATAC-seq data, parallel processing is needed to limit runtime (Figure 6d). Runtime increases decidedly when applying SEMITONES to larger numbers of features and reference cells, or combinations thereof (see supplementary Figure 12, Additional File 1). Currently, the main bottleneck lies in the memory demand for large numbers of reference cells, because the similarity matrix is dense and of the dimension |cells| × |reference cells|. Therefore, it is advisable to use multiple cores when using a large number of features for very sparse data, and submitting individual jobs for subsets of reference cells when applying semitones to large numbers of reference cells.

**Figure 6:**
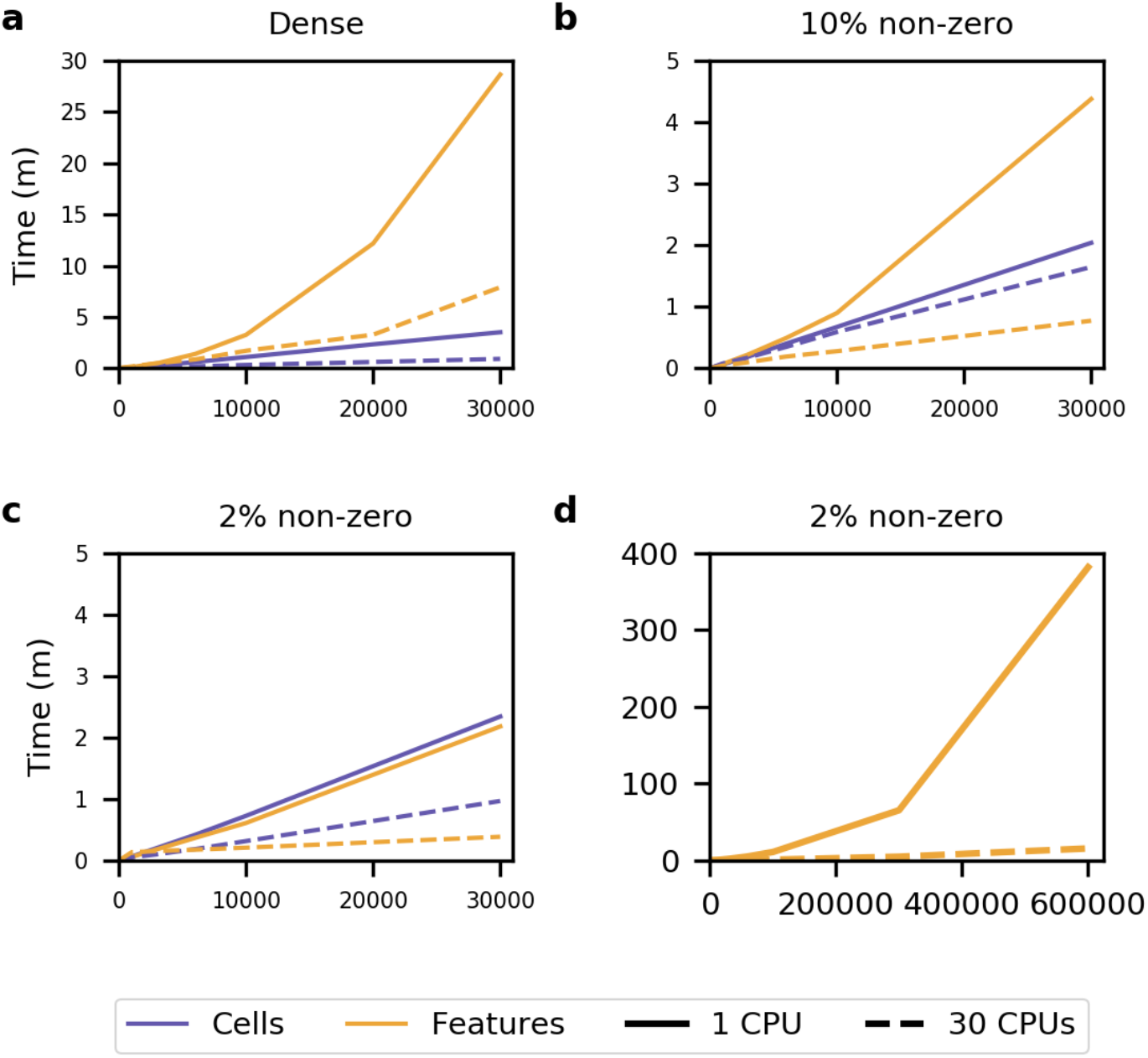
Runtime scalability of SEMITONES. The runtime of SEMITONES when applied to a) data with all non-zero values, b) data with a density representative of scRNA-seq data (10% non-zero values), and c) a data density representative of scATAC-seq data (2% non-zero values) for a maximum of 30,000 features. d) The runtime of SEMITONES when applied to data with 2% non-zero values with a maximum of 600,000 features, which isrepresentative for a large scATAC-seq data set that was not filtered for highly variably/commonly accessible regions. The purple lines show the runtime with respect to the number of reference cells and the orange lines show the runtime with respect to the number of features. The dotted lines represent the runtime when parallelizing over 30 CPUs and the solid lines show the runtime when using 1 CPU. All results were obtained for a dataset containing 100,000 rows, simulating a data set of 100,000 cells in total.

## Discussion

We present SEMITONES; a tool for the *de novo* identification of informative features in single-cell omics data. We illustrate that SEMITONES identifies marker genes and regulators of cell identity without first clustering the cells. This way, we aim to mitigate the propagation of errors or biases from cluster assignments. Additionally, we show that SEMITONES is an effective alternative to highly variable genes for feature selection in scRNA-seq preprocessing.

Here, SEMITONES identifies a smaller number of genes that captures the same biological diversity as a larger number of HVGs. Lastly, we show that SEMITONES can also be readily applied to the identification of relevant peaks from scATAC-seq data. In short, SEMITONES is a flexible tool aiding the identification of biologically relevant features from single-cell omics data.

SEMITONES accurately retrieves marker genes of cell identity. Since reference cell annotations based on these markers largely overlap with the published annotations [5], we conclude that SEMITONES accurately retrieves cell identity specific markers, and propose highly enriched genes for which we did not find literature evidence, like *PADI4* in immature classical monocytes, as putative novel markers. These highly specific marker genes also annotated subpopulations of cells that are otherwise highly similar, like monocytes and T cells. The use of an RBF-kernel to describe cell similarities enables the identification across a broad range of specificity because we can use the parameter γ to define the cell neighbourhood range for which we identify informative genes (see supplementary Figure 5, Additional File 1). By performing regression to these similarities, we remove the need to assign cells to groups of the same identity but instead allow cells to be part of multiple cell neighbourhoods. This way, we identify marker genes along the haematopoietic axis, as was illustrated for the B cell lineage. On the other hand, the dependence on the similarity metric is a potential limitation of SEMITONES, especially when no prior knowledge is available to evaluate the adequacy of the similarity metric. Importantly, an adequate similarity metric is essential for the data-driven selection of a set of reference cells that is representative of the biological cell diversity. Ultimately, given an accurate similarity metric, SEMITONES identifies highly specific markers of cell identity, illustrated here for the haematopoietic axis. We also explore using highly enriched genes, identified by SEMITONES, as an alternative to using HVGs. Cell identity separation and localization of marker gene expression in the 2D UMAP space improves when using highly enriched genes compared to HVGs. Thus, we conclude that n highly enriched genes capture more biological variability than n HVGs. This is of particular interest in light of recent developments in targeted scRNA-seq, for which the need to select a set of a few hundred genes arises that contain sufficient transcriptomic information regarding the biological system. The adequate performance while using the cosine similarity over the top 50 principal components instead of an RBF-kernel over 25 UMAP dimensions illustrates that SEMITONES also identifies informative genes when using a non-specialized similarity metric. However, the performance of SEMITONES will depend on the provided reference cells since SEMITONES only identifies highly enriched genes in reference-cell neighbourhoods. From simulations (see supplementary Figure 4, Additional File 1) we conclude that reference cell sets obtained using suboptimal similarity metric-embedding combinations do not represent all cell identities. In this case, the highly enriched genes will only capture the variability for cell identities present in the reference cells. Based on the same simulations, we recommend using the Euclidean distance over a reasonable number (e.g. 50) of principal components if using the SEMITONES data-driven reference cell selection. Besides marker gene identification, SEMITONES can be used to construct co-enrichment graphs. These co-enrichment graphs revealed several interactions indicative of mechanistic aspects of gene regulation and cellular function. For example, high co-enrichment of *CD8A, CD3E*, and *CD3D* in CD8^+^ T lymphocytes is substantiated by the role of *CD8A* as a chaperone to the CD3-complex (Figure 3a). Highly central nodes (i.e. genes) in these networks represent putative regulators of cell identity, even if they are not individually enriched in a cell neighbourhood, as seen for *S100A4* gene in T_h_17 cells (Figure 3b, [35]).

The SEMITONES workflow is readily transferable to the identification of marker peaks from scATAC-seq [peak x cell] matrices. Based on GO term enrichment of associated genes and enrichment for known transcription factor binding motifs, we conclude that SEMITONES retrieves biologically relevant peaks. Additionally, we identify biological signatures that we did not unveil in the marker genes. This illustrates the benefit of analysing scATAC-seq data independently of scRNA-seq data, although we cautiously note that this observation may be a result of differences in the cell identities represented by the reference cells. Besides using the enriched peaks to annotate cell identities, we also use significantly enriched peaks to identify global patterns of chromatin accessibility and *cis*-regulatory regions. Most notably, we find that selectively inaccessible regions are most likely promoters and selectively accessible regions are most likely enhancers, once more indicating the higher cell type-specificity of enhancers. However, these results are limited to a small number of reference cells and were not subjected to rigorous statistical analysis.

## Conclusion

SEMITONES is a diverse and flexible tool for the identification of informative features from single-cell omics data, readily applicable to expression and chromatin-related data. Its possible limitations include the need for an adequate cell similarity metric and a set of reference cells that is representative of the cell population. Therefore, in future research, we will explore deterministic approaches and the use of geometric sketching [44] to select an optimal set of reference cells. Additionally, we aim to improve the run time for many features and the memory demand for large numbers of reference cells. Namely, the appli-cation of SEMITONES to large numbers of reference cells currently requires the user to perform computations for subsets of reference cells due to limitations in the multiprocessing setup. On a biological level, we will explore the integration of scRNA-seq and scATAC-seq data on the cell level. As such, we aim to keep SEMITONES up to date as single-cell data grows and diversifies, to aid the elucidation of regulatory mechanisms underlying the acquisition of cell identity in health and disease.

## Methods

### Reference cell selection

In this article, we use two reference cell selection methods: an automated data-driven cell selection method, and manual selection of a set of reference cells from a 2D cell embedding. For the manual selection of cells from any 2D embedding, we provide a figure widget implementation. The data-driven cell selection method is presented in Algorithm 1 and described below.

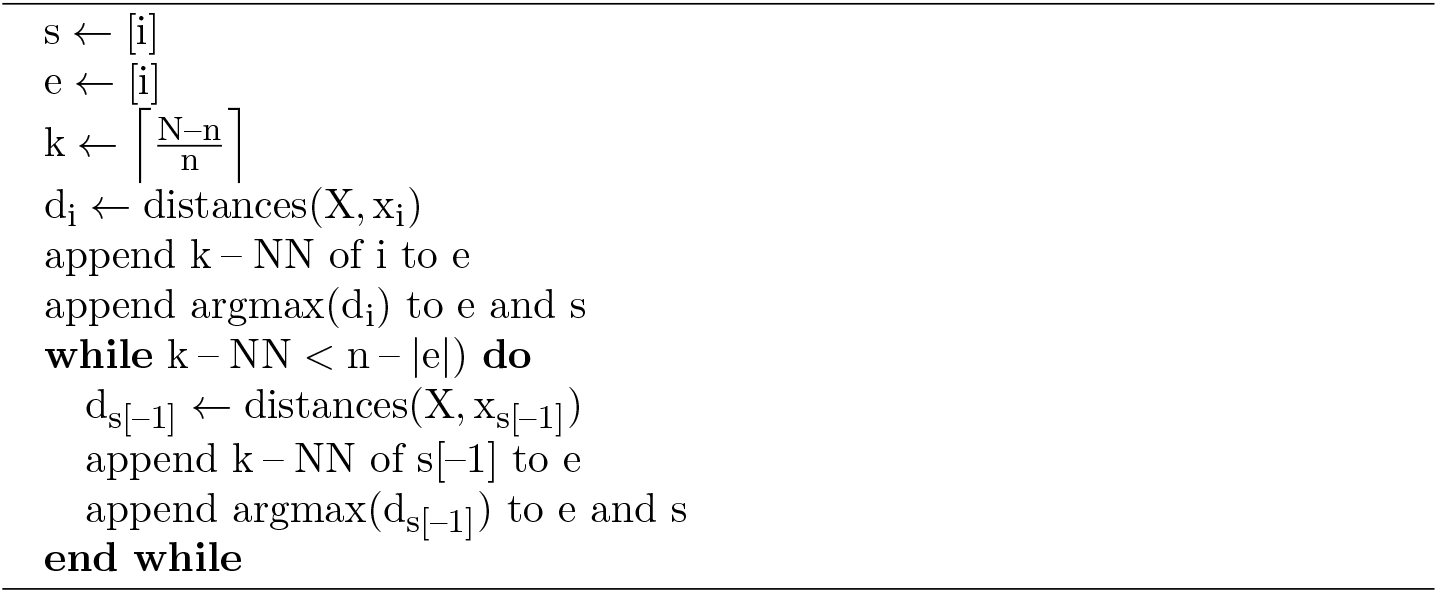

In addition to the methods applied in this study, we provide the options to use the sklearn k-means++ implementation and a fixed-grid search. In the fixed-grid search, a lattice graph of a user-defined size n × n is placed over the 2D embedding of single cells, as illustrated in Supplementary Figure 13, Additional File 1. Then, the cells closest to the intersections of the horizontal and vertical grid lines are selected. The method avoids selecting a disproportionate number of cells at the edge of the 2D-embedding by putting a constraint on the minimum distance between each pair of selected cells. The implementations of these methods can be found in the cell selection module of SEMITONES.

### Enrichment scoring

Given a set of reference cells (Figure 1a), we identify informative features in these cells. From the idea of an informative feature as being only expressed or absent in similar cells, we can derive the formal definition that informative features harbour a strong linear relationship with the similarity to the reference cell (Figure 1b).

In SEMITONES, we infer informative features using a simple linear regression framework (Equation 1). Here, y_c_ is a vector representing each cell by its similarity to some reference cell c using any suitable metric. As example, we use an RBF-kernel with γ = 8 × 10^−1^ over a multidimensional UMAP embedding in applications for marker selection. For applications to feature selection, we use the cosine similarity over the top n principal components. The vector x_f_ represents the value of the feature g in each cell. When applying the method to scRNA-seq data, this feature vector x_f_ contains the gene expression level in each cell. For applications to scATAC-seq data, x_f_ is a binary feature vector indicating whether the chromatin at a certain location in a cell is accessible (1) or not (0). The regression coefficient β_c,f_, which is estimated using the ordinary least squares method, describes the strength of the linear relationship between y_c_ and x_f_. Thus, the value of β_c,f_ can be interpreted as a score of the enrichment of some feature f in some reference cell c. High positive enrichment scores will be obtained for features which are only observed in cells similar to some cell c (Figure 1b). Inversely, low negative scores will be obtained for features which are only observed in cells which are dissimilar to some cell c (Figure 1b).

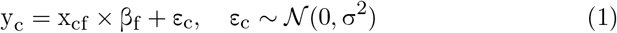

In addition to single feature enrichment scores, one can also opt to calculate enrichment scores for sets of features. In this case, x_f_ is a vector representing the combined values of all features in the set. In the case of continuous feature values, like in gene expression values in scRNA-seq, we provide four different approaches to representing a set of features in a single vector. The first approach is the multiplication of the vectors, like an interaction term in multiple regression. The second and third approaches are to select the lowest or highest expression value of the features in a set as the representative expression value, respectively. Lastly, the fourth approach is to take the median expression of the features in the set as the representative value. For binary feature vectors, like in scATAC-seq, we can also readily take the median feature values to present the feature set expression vector. Additionally, we implement the strategy of annotation a feature set as present (1) if one or all of the features in a set are present and absent (0) if none of the features is present.

The pairwise feature set enrichment scores can be used to construct coenrichment graphs, where vertices (i.e. features) are connected by edges that are weighted by the feature set enrichment scores (Figure 1d). To improve interpretability we then infer the maximum spanning tree of these graphs, in which all vertices are connected using the least number of edges with a maximum total weight, using networkx (v2.4) [45]. The current flow betweenness centrality measure is used as a measure of the importance of a feature in the co-enrichment network. Visualization of graphs is performed in Netwulf [46].

### Significance testing

The null distribution for significance testing is obtained by repeating the scoring procedure using n times permuted feature vectors. Due to the permutation of the feature vectors, the feature values are randomized while still resembling the original data. Significance is declared at a user-defined number of standard deviations (n_σ_) away from the mean of this null distribution (Figure 1c). Here, we always use n = 256.

### Data processing

The practical use of SEMITONES is illustrated by its application to healthy haematopoiesis scRNA-seq and scATAC-seq data published by [5]. The scRNA-seq count matrices were obtained from the GEO database (GSE139369, accessed February 28 2020). The scATAC-seq count matrix was downloaded from the GitHub page linked to the original data publication ([47], accessed on 3 March 2020).

The scRNA-seq data covers a total of 35582 cells obtained from six different samples, including two samples of CD34^+^ enriched BMMCs, two samples of non-enriched BMMCs and two samples of PBMCs. First, we removed any cells for which the ratio between the number of genes expressed over the count-depth is greater than or equal to 0.3. Next, we performed scran deconvolution normalization using the computeSumFactors function using clusters obtained from the quickCluster function [48, 49]. The normalized counts were log-transformed using an alternative pseudo-count as proposed by Lun et al. (2018) [50]. Inspection of the count depth of cells in a 2D UMAP embedding (computed over the top 10 principal components) revealed a cluster of cells with low count depth in one of the CD34^+^ samples, which was removed. This leaves a total of 35156 cells.

The (non-normalized) count data from all cells that passed quality control were combined, and scran deconvolution normalization was performed on the combined data. The data were log-transformed using an alternative pseudocount [50] for use in enrichment scoring. We then performed latent semantic indexing for the reduction of the normalized count data to a 50-dimensional embedding. A 2D and 25D uniform manifold approximation and projection (UMAP, 30 neighbours and a minimum distance of 0.3) over the LSI space were computed for visualization and similarity calculations, respectively.

The scATAC-seq data contains a total of 35038 DC34^+^ enriched BMMC, non-enriched BMMC, and PBMC cells. We performed quality control on the combined data as follows. Cells were removed if their peak depth exceeds 200,000 or more than 60,000 peaks were called in this cell, and peaks were removed if their count exceeds 40,000, leaving 35022 cells. Next, we binarized the peak by cell-matrix and perform LSI to reduce the feature space to 50 dimensions. We computed a 2D and 35D UMAP (50 neighbours, minimum distance of 0.5) over the 50-dimensional space for visualization and similarity calculations, respectively.

### Evaluation references

For the annotation of the scRNA-seq reference cells we look at the top 10 most highly enriched genes (according to SEMITONES). The Human Blood Atlas [9], with a special focus on the Monaco scaled dataset [51], served as a primary reference. Additional markers were obtained from the literature (see Supplementary Table 2, Additional File 3). The STRING (v11.0) database ([29], [52]) was used for qualitative evaluation of co-enrichment graphs. The assessment of SEMITONES as a method for feature selection in scRNA-seq was performed in comparison to the retrieval of highly variable genes as implemented in Scanpy (v1.4.5) [39]. For the annotation of scATAC-seq reference cells, we obtain GO-term enrichments and associated genes for significantly enriched peaks using GREAT (v4.0.4) [40]. We provide all peaks in the clean scATAC-seq data as a background set and select the “basal plus extension” association rule to characterize the regulatory domain. According to this association rule, the proximal domain is 5 kilobases upstream and 1 kilobase downstream of the transcription start site (TSS), and the distal domain is defined as up to 1000 kilobases from the TSS. *cis*-Regulatory element annotations were obtained from HOMER (v10.4) (promoter, exon, 5’ UTR, 3’ UTR, intronic, intergenic, transcription termination side) and the permissive enhancer annotations in FANTOM 5 phase 2.6. In HOMER (v10.4), a region is annotated as a promoter of it lies within −1000 and +100 base pairs from the TSS as annotated in RefSeq. Motif enrichment of known transcript factor (TF) binding motifs was performed using the findMotifsGenome function from HOMER (v10.4). We consider motifs enriched if their (Benjamini) q-value < 0.01.

### Scalability of run time

The run time scalability was assessed for different numbers of cells, numbers of features, data densities, and the number of core processing units (CPUs). Random data sets with 100%, 10%, or 2% nonzero values were constructed. The decision for 10% and 2% nonzero values were based on the sparsity character of the data used in the application example. In all experiments, the total number of cells was set to be 100,000. Run time was compared between computations using one CPU and 30 CPUs. Parallelization over rows or columns was selected based on whether the number of rows or columns was greater, respectively.

### Availability of data and materials

The datasets supporting the conclusions of this article are available from public sources. The healthy haematopoiesis scRNA-seq dataset was downloaded from the Gene Expression Omnibus (GSE139369). The healthy scATAC-seq dataset was downloaded from GitHub, accessed on 3 March 2020 [47, 5]. The SEMITONES software is freely available from GitHub [53] under the GPL-3.0 license. The scripts and notebooks used for data processing and analyses are published on GitHub [54].

## Author’s contributions

AHCV, SM, and UO conceived the project. AHCV implemented and evaluated the method. UO and SM guided the implementation of the method. UO guided the evaluation of the method. AHCV wrote the draft manuscript and UO and SM suggested revisions. All authors approved the final manuscript.

## Additional Files

**Additional file 1 — Supplemental Figures**

This file (.pdf) contains all supplemental figures that were referenced within the main text.

**Additional file 2 — Table S1**

Table S1 (.csv) lists the top 10 most highly enriched genes for all data-driven selected reference cells.

**Additional file 3 — Table S2**

Table S2 (.csv) lists the marker genes used for the annotation of reference cells based on the top 10 most highly enriched genes.

**Additional file 4 — Table S3**

Table S3 (.csv) lists the most central nodes in the co-enrichment graphs constructed using enrichment scores for interaction vectors, maximum-value vectors, medianvalue vectors, minimum-value vectors.

## References

[1] Luecken MD, Theis FJ. Current best practices in single-cell RNA-seq analysis: a tutorial. Molecular Systems Biology. 2019;15(6):e8746.

[2] Townes FW, Hicks SC, Aryee MJ, Irizarry RA. Feature selection and dimension reduction for single-cell RNA-seq based on multinomial model. Genome Biology. 2019;20:295.

[3] Lähnemann D, Köster J, Szczurek E, McCarthy D, Hicks S, Robinson M, et al. Eleven grand challenges in single-cell data science. Genome Biology. 2020;21.

[4] Farrell JA, Wang Y, Riesenfeld SJ, Shekhar K, Regev A, Schier AF. Single-cell reconstruction of developmental trajectories during zebrafish embryogenesis. Science. 2018;360(6392).

[5] Granja JM, Klemm S, McGinnes LM, Kathiria AS, Mezger A, Corces MR, et al. Single-cell multiomic analysis identifies regulatory programs in mixed-phenotype acute leukemia. Nature Biotechnology. 2019;37:1458–1465.

[6] Velten L, Haas S, Raffel S, Blaszkiewicz S, Islam S, Hennig BP, et al. Human haematopoietic stem cell lineage commitment is a continuous process. Nature cell biology. 2017;19:271–281.

[7] Laabi Y, Gras MP, Brouet JC, Berger R, Larsen CJ, Tsapis A. The BCMA gene, preferentially expressed during B lymphoid maturation, is bidirectionally transcribed. Nucleic Acids Research. 1994;22(7):1147–1154.

[8] Smith EL, Harrington K, Staehr M, Masakayan R, Jones J, Long TJ, et al. GPRC5D is a target for the immunotherapy of multiple myeloma with rationally designed CAR T cells. Science Translational Medicine. 2019;11(485).

[9] Uhlén M, Karlsson MJ, Zhong W, Tebani A, Pou C, Mikes J, et al. A genomewide transcriptomic analysis of protein-coding genes in human blood cells. Science. 2019;366.

[10] Zheng S, Papalexi E, Butler A, Stephenson W, Satija R. Molecular transitions in early progenitors during human cord blood hematopoiesis. Molecular Systems Biology. 2018;14(3):e8041.

[11] Toren AG, Bielorai B, Jacob-Hirsch J, Fisher T, Kreiser D, Moran O, et al. CD133-positive hematopoietic stem cell “stemness” genes contain many genes mutated or abnormally expressed in leukemia. Stem cells. 2005;23 8:1142–53.

[12] He X, Gonzalez V, Tsang AH, Thompson JA, Tsang TC, Harris DT. Differential gene expression profiling of CD34+ CD133+ umbilical cord blood hematopoietic stem progenitor cells. Stem cells and development. 2005;14 2:188–98.

[13] Huang H, Sikora MJ, Islam S, Chowdhury RR, Chien Yh, Scriba TJ, et al. Select sequencing of clonally expanded CD8+ T cells reveals limits to clonal expansion. Proceedings of the National Academy of Sciences of the United States of America. 2019;116:8995–9001.

[14] Viemann D, Strey A, Janning A, Jurk K, Klimmek K, Vogl T, et al. Myeloid-related proteins 8 and 14 induce a specific inflammatory response in human microvascular endothelial cells. Blood. 2005;105(7):2955–62.

[15] Roth J, Goebeler M, Wrocklage V, van den Bos C, Sorg C. Expression of the calcium-binding proteins MRP8 and MRP14 in monocytes is regulated by a calcium-induced suppressor mechanism. The Biochemical journal. 1994;301 (Pt 3).

[16] Odink K, Cerletti N, Br?ggen J, Clerc RG, Tarcsay L, Zwadlo G, et al. Two calcium-binding proteins in infiltrate macrophages of rheumatoid arthritis. Nature. 1987;330(6143):80–82.

[17] Zhao F, Hoechst B, Austin D, Gamrekelashvili J, Fioravanti S, Manns MP, et al. S100A9 a new marker for monocytic human myeloid derived supressor cells. Immunology. 2012;136:176–183.

[18] Hofer TP, Zawada AM, Frankenberger M, Skokann K, Satzl AA, Gesierich W, et al. slan-defined subsets of CD16-positive monocytes: impact of granulomatous inflammation and M-CSF receptor mutation. Blood. 2015;126(24):2601–2610.

[19] Rothenberg EV. Transcriptional Control of Early T and B Cell Developmental Choices. Annual Review of Immunology. 2014;32(1):283–321.

[20] Kohn L, Hao QL, Sasidharan R, Parekh C, Ge S, Zhu Y, et al. Lymphoid Priming in Human Bone Marrow Begins Prior to CD10 Expression with UpRegulation of L-selectin. Nature immunology. 2012 09;13:963–71.

[21] Kartal-Kaess M, Cimmino L, Infantino S, Yabas M, Zhang JG, Tuohey L, et al. RNAi Screening Identifies A Novel Role for A-Kinase Anchoring Protein 12 (AKAP12) in B Cell Development and Function. Blood. 2012;120(21):855–855.

[22] O’Leary NA, Wright MW, Brister JR, Ciufo S, Haddad D, McVeigh R, et al. Reference sequence (RefSeq) database at NCBI: current status, taxonomic expansion, and functional annotation. Nucleic Acids Research. 2015 11;44(D1):D733–D745.

[23] Zandi S, Månsson R, Tsapogas P, Zetterblad J, Bryder D, Sigvardsson M. EBF1 Is Essential for B-Lineage Priming and Establishment of a Transcription Factor Network in Common Lymphoid Progenitors1. The Journal of Immunology. 2008;181:3364–3372.

[24] Clark MR, Mandal M, Ochiai K, Singh H. Orchestrating B cell lymphopoiesis through interplay of IL-7 receptor and pre-B cell receptor signalling. Nature reviews Immunology. 2014;14(2):69–80.

[25] Suryani S, Fulcher DA, Santner-Nanan B, Nanan R, Wong M, Shaw PJ, et al. Differential expression of CD21 identifies developmentally and functionally distinct subsets of human transitional B cells. Blood. 2010;115 3:519–29.

[26] Bigot J, Pilon C, Matignon M, Grondin C, Leibler C, Aissat A, et al. Transcriptomic Signature of the CD24hi CD38hi Transitional B Cells Associated With an Immunoregulatory Phenotype in Renal Transplant Recipients. American journal of transplantation: official journal of the American Society of Transplantation and the American Society of Transplant Surgeons. 2016;16(12):3430–3442.

[27] Said JW, Hoyer KK, French SW, Rosenfelt L, Garcia-Lloret MI, Koh PJ, et al. TCL1 Oncogene Expression in B Cell Subsets from Lymphoid Hyperplasia and Distinct Classes of B Cell Lymphoma. Laboratory Investigation. 2001;81:555–564.

[28] Polyak MJ, Li H, Shariat N, Deans JP. CD20 Homo-oligomers Physically Associate with the B Cell Antigen Receptor. Journal of Biological Chemistry. 2008;283:18545–18552.

[29] Szklarczyk D, Gable AL, Lyon D, Junge A, Wyder S, Huerta-Cepas J, et al. STRING v11: protein–protein association networks with increased coverage, supporting functional discovery in genome-wide experimental datasets. Nucleic Acids Research. 2019;47:D607–D613.

[30] Wang LD, Clark MR. B-cell antigen-receptor signalling in lymphocyte development. Immunology. 2003;1104:411–20.

[31] Artyomov MN, Lis M, Devadas S, Davis MM, Chakraborty AK. CD4 and CD8 binding to MHC molecules primarily acts to enhance Lck delivery. Proceedings of the National Academy of Sciences. 2010;107:16916–16921.

[32] Zawada AM, Rogacev KS, Rotter B, Winter P, Marell RR, Fliser D, et al. SuperSAGE evidence for CD14++CD16+ monocytes as a third monocyte subset. Blood. 2011;118 12:e50–61.

[33] Vassel L, Beauchemin H, Lemsaddek W, Krongold J, Trudel M, Möröy T. Growth factor independence 1b (gfi1b) is important for the maturation of erythroid cells and the regulation of embryonic globin expression. PLoS One. 2014;9(5):e96636.

[34] da Cunha AF, Brugnerotto AF, Duarte ASS, Lanaro C, Costa GGL, Saad STO, et al. Global gene expression reveals a set of new genes involved in the modification of cells during erythroid differentiation. Cell proliferation. 2010;43:297–309.

[35] Brisslert M, Bian L, Svensson MND, Santos RMF, Jonsson IM, Barsukov IL, et al. S100A4 regulates the Src-tyrosine kinase dependent differentiation of Th17 cells in rheumatoid arthritis. Biochimica et biophysica acta. 2014;184211:2049–59.

[36] Zemmour D, Zilionis R, Kiner E, Klein AM, Mathis D, Benoist C. Single-cell gene expression reveals a landscape of regulatory T cell phenotypes shaped by the TCR. Nature immunology. 2018;19:291–301.

[37] Kihm AJ, Kong Y, Hong W, Russell JE, Rouda S, Adachi K, et al. An abundant erythroid protein that stabilizes free alpha-haemoglobin. Nature. 2002;417:758–763.

[38] Magor GW, Tallack MR, Gillinder KR, Bell CC, McCallum N, Williams BA, et al. KLF1-null neonates display hydrops fetalis and a deranged erythroid transcriptome. Blood. 2015;125 15:2405–2417.

[39] Wolf F, Angerer P, Theis F. SCANPY: large-scale single-cell gene expression data analysis. Genome Biology. 2018;19:15.

[40] McLean CY, Bristor D, Hiller M, Clarke SL, Schaar BT, Lowe CB, et al. GREAT improves functional interpretation of cis-regulatory regions. Nature Biotechnology. 2010;28(5):495–501.

[41] Wang H, Lee CH, Qi C, Tailor P, Feng J, Abbasi S, et al. IRF8 regulates B-cell lineage specification, commitment, and differentiation. Blood. 2008;112 10:4028–38.

[42] Ho IC, Tai TS, Pai SY. GATA3 and the T-cell lineage: essential functions before and after T-helper-2-cell differentiation. Nature Reviews Immunology. 2009;9:125–135.

[43] Natarajan A, Yardimci GG, Sheffield NC, Crawford GE, Ohler U. Predicting cell-type-specific gene expression from regions of open chromatin. Genome Research. 2012;22:1711–1722.

[44] Do VH, Elbassioni K, Canzar S. Sphetcher: Spherical Thresholding Improves Sketching of Single-Cell Transcriptomic Heterogeneity. iScience. 2020;23(6):101126.

[45] Hagberg AA, Schult DA, Swart PJ. Exploring Network Structure, Dynamics, and Function using NetworkX. Proceedings of the 7th Python in Science Conference. 2008:11–15.

[46] Aslak U, Maier BF. Netwulf: Interactive visualization of networks in Python. Journal of Open Source Software. 2019;4(42):1425. Available from: https://doi.org/10.21105/joss.01425.

[47] Granja JM. GreenleafLab MPAL single-cell 2019 GitHub;. Available from: https://github.com/GreenleafLab/MPAL-Single-Cell-2019. Accessed on 3 March 2020.

[48] Lun ATL, McCarthy DJ, Marioni JC. A step-by-step workflow for low-level analysis of single-cell RNA-seq data with Bioconductor. F1000Res. 2016;5:2122.

[49] Lun ATL, Bach K, Marioni JC. Pooling across cells to normalize single-cell RNA sequencing data with many zero counts. Genome Biology. 2016;17:75.

[50] Lun A. Overcoming systematic errors caused by log-transformation of normalized single-cell RNA sequencing data. bioRxiv. 2018.

[51] Monaco G, Lee B, Xu W, Mustafah S, Hwang YY, Carre C, et al. RNA-Seq Signatures Normalized by mRNA Abundance Allow Absolute Deconvolution of Human Immune Cell Types. Cell Reports. 2019;26:1627–1640.e7.

[52] 2020 SC. STRING Database;. Available from: https://string-db.org.

[53] Vlot AHC. SEMITONES;. Available from: https://github.com/ohlerlab/SEMITONES.

[54] Vlot AHC. SEMITONES;. Available from: https://github.com/ohlerlab/SEMITONES_paper.

