## Additional file 1 - Supplemental Figures for "SEMITONES: Single-cEll Marker IdentificaTiON by Enrichment Scoring"

### Results

#### SEMITONES identifies marker genes

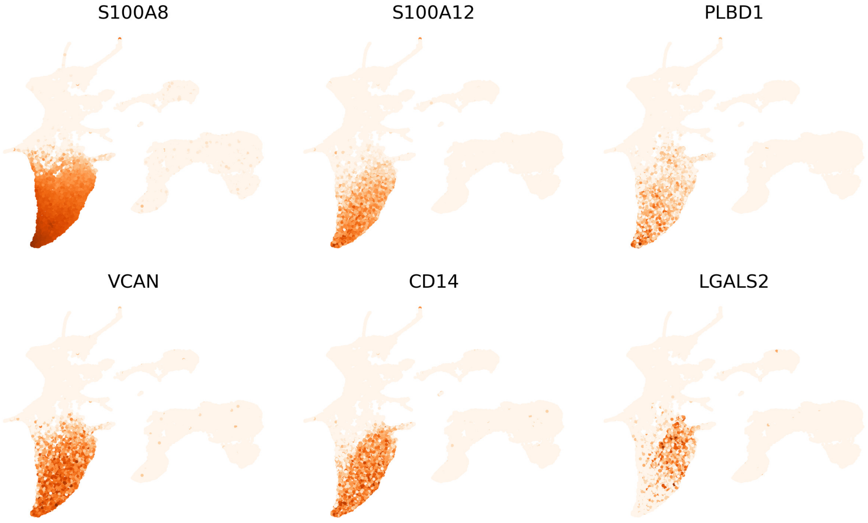


**Figure S1:** Markers of immature calssical monocytes (top row) and intermediate and classical monocytes (bottom row).

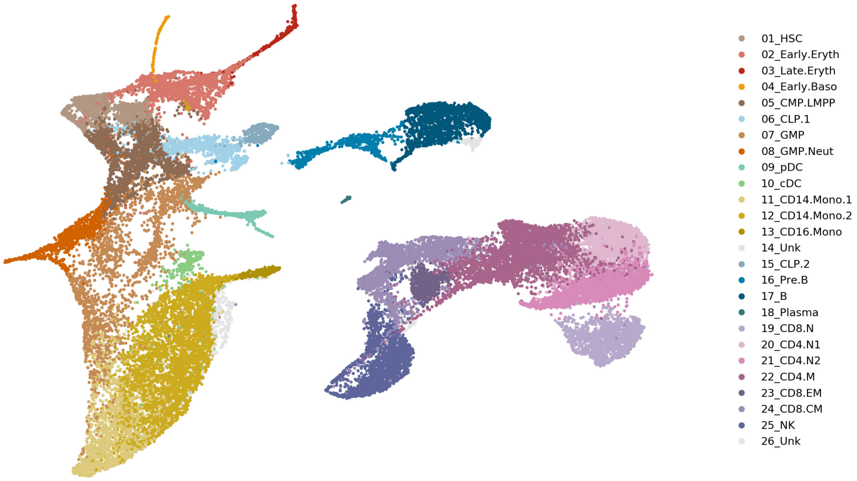


**Figure S2:** Cluster-based biological cell type annotations as taken from [1].

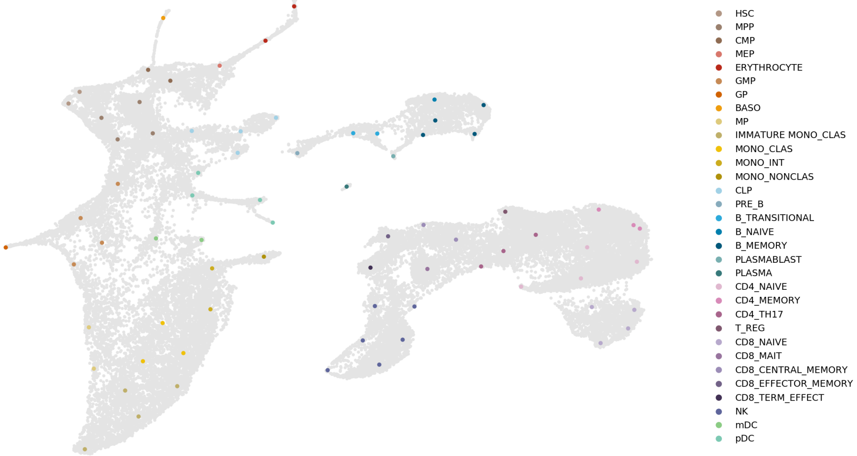


**Figure S3:** The annotation of manually selected reference cells from scRNA-seq data.

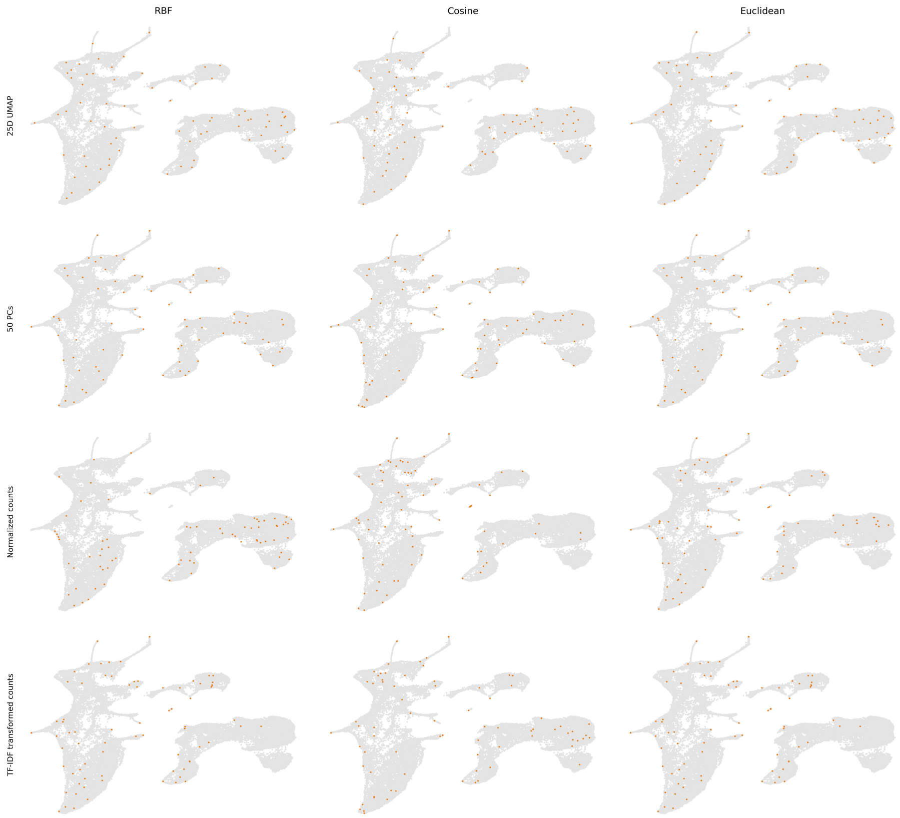


**Figure S4:** Data-driven cell selection using different embeddings (rows) and distance metrics (columns).

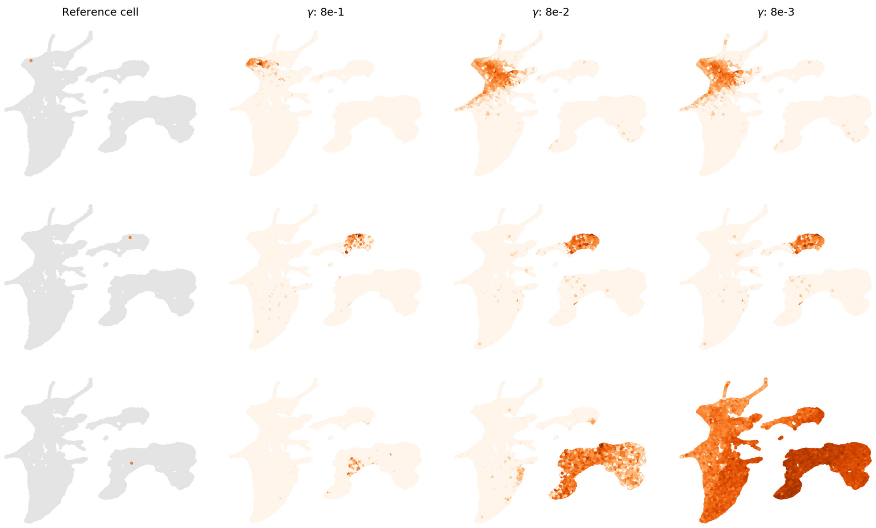


**Figure S5:** The influence of the radius of influence when using the RBF-kernel as a similarity metric. A higher value of $\gamma$ is proportional to a larger radius of influence in the RBF-kernel.

#### SEMITONES identifies transcriptional regulators

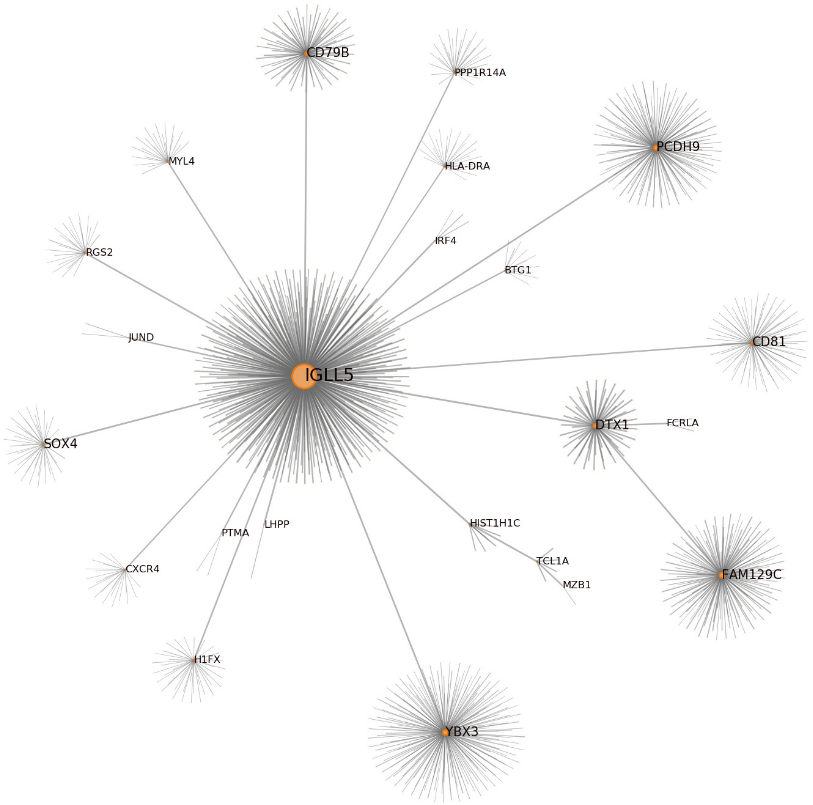


**Figure S6:** The interaction co-enrichment graph for the transitional B cell neighbourhood.

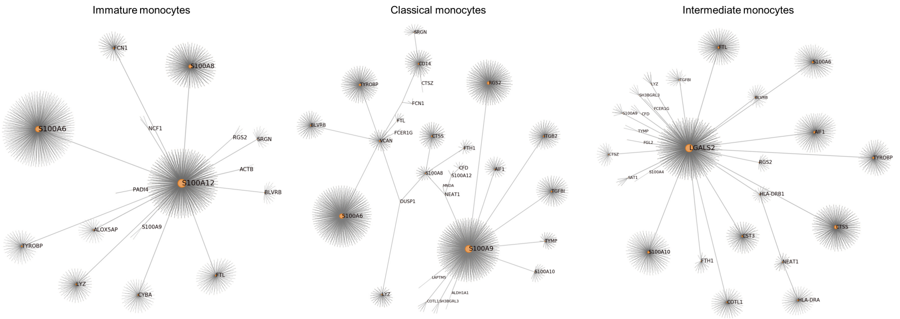


**Figure S7:** The interaction co-enrichment graphs for distinct, but highly similar, monocyte populations.

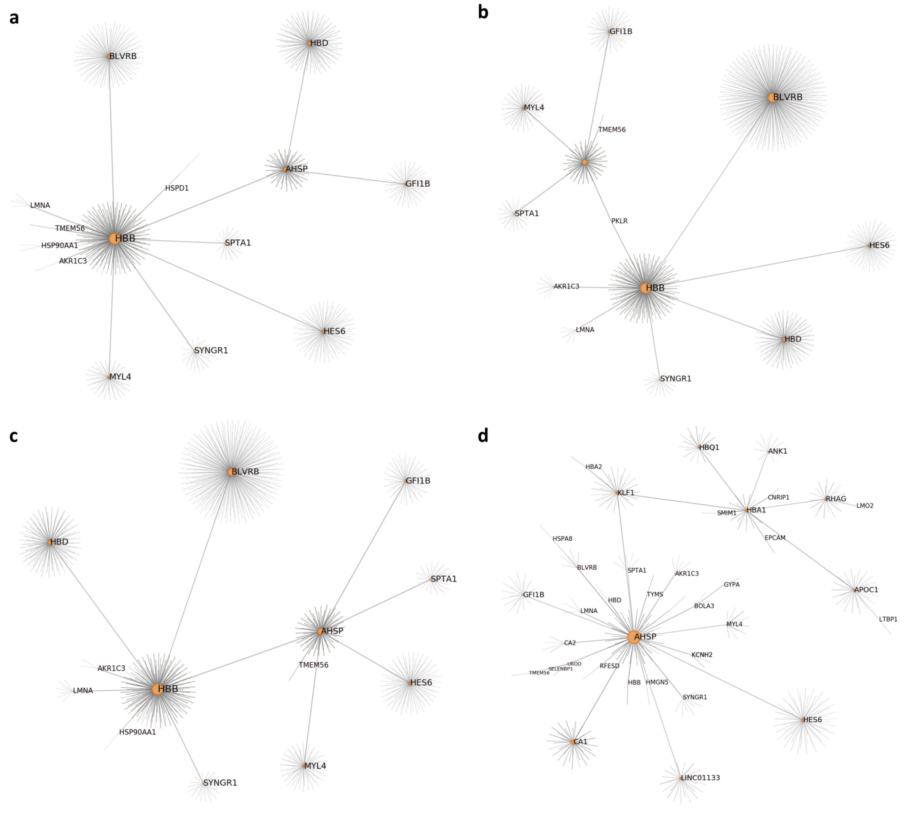


**Figure S8:** Erythrocyte a) interaction co-enrichment graph, b) maximum-value co-enrichment graph, c) median-value co-enrichment graph, d) minimum-value co-enrichment

#### SEMITONES for feature selection

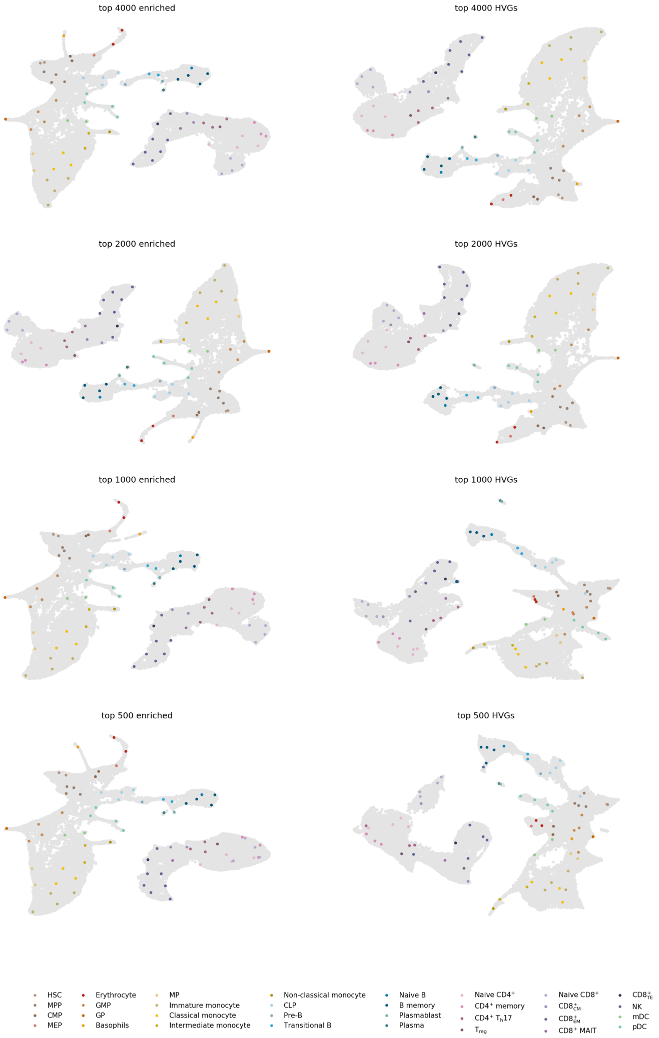


**Figure S9:** UMAPs obtained using different numbers of top enriched (left) or highly variable genes (right).

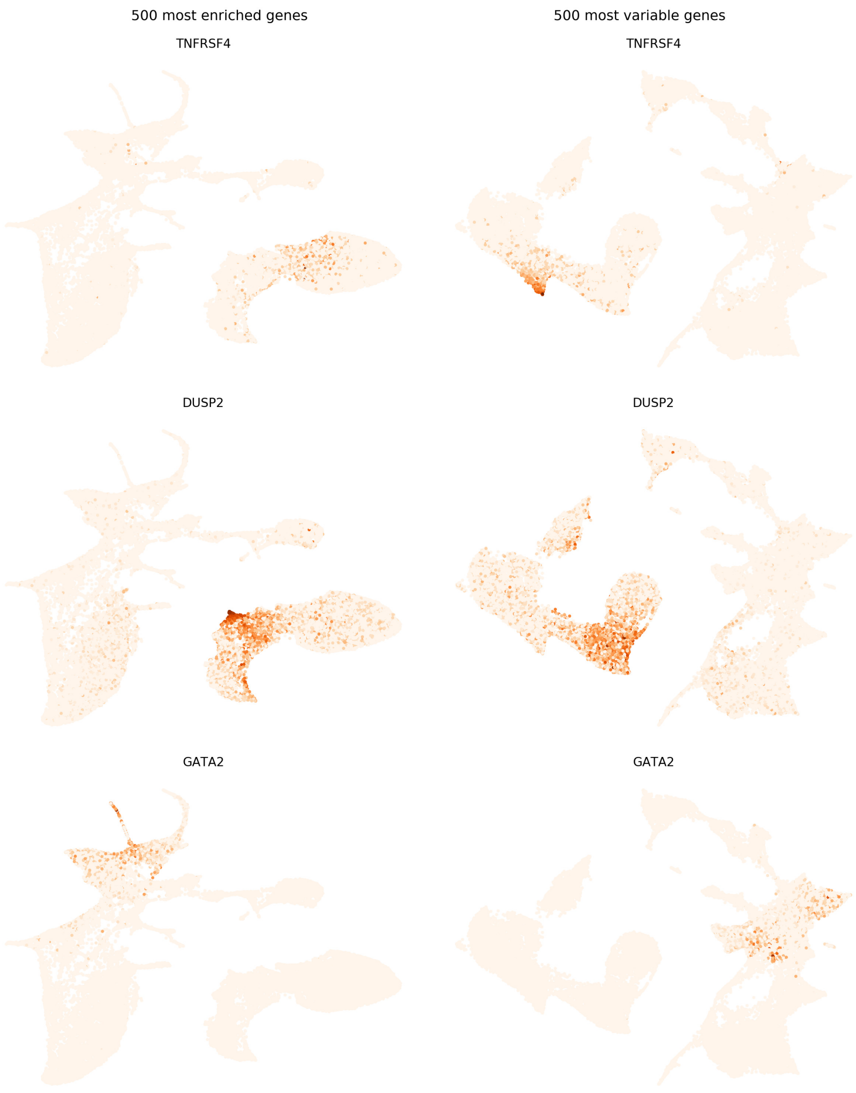


**Figure S10:** The expression levels of marker genes in 2D UMAPs computed using 500 most enriched genes (left) or 500 most variable genes (right).

#### SEMITONES identifies cell-specific cis-regulatory elements

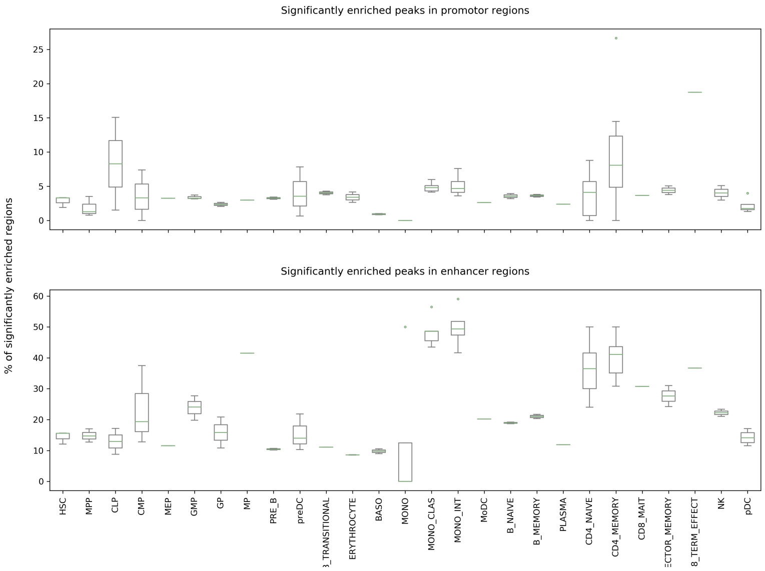


**Figure S11:** Percentage of signficantly positively enriched peaks that fall in promoter (top panel) or enhancer (bottom panel) regions.

### Scalability of SEMITONES

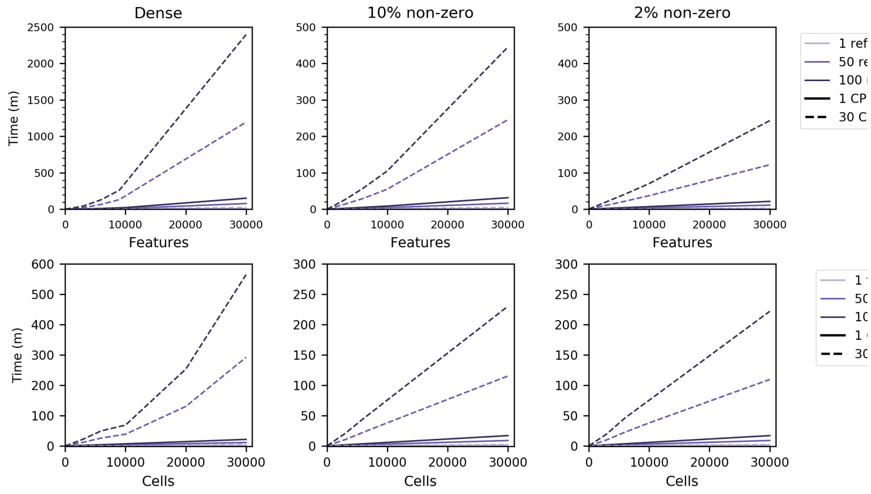


**Figure S12:** The run time of SEMITONES for 1-30,000 features or reference cells (top row) and 1-100 features or reference cells (bottom row).

### Methods

#### Reference cell selection

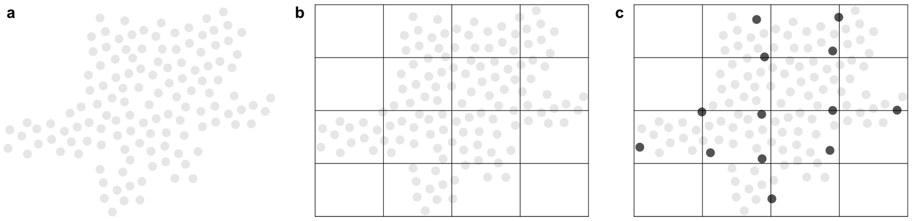


**Figure S13:** Fixed-grid cell selection. Given a 2D cell embedding (a), we fit a lattice graph of size $n \times n$ (b) and then select cells closest to the intersections of the horizontal and vertical grid lines (c).

### References

[1] Granja JM, Klemm S, McGinnes LM, Kathiria AS, Mezger A, Corces MR, et al. Single-cell multiomic analysis identifies regulatory programs in mixed-phenotype acute leukemia. Nature Biotechnology. 2019;37:1458–1465.
